# Description of new micro-colonial fungi species *Neophaeococcomyces mojavensis*, *Coniosporium tulheliwenetii, and Taxawa tesnikishii* cultured from biological soil crusts

**DOI:** 10.1101/2024.06.12.598762

**Authors:** Tania Kurbessoian, Sarah A. Ahmed, Yu Quan, Sybren de Hoog, Jason E. Stajich

## Abstract

Black yeasts and relatives comprise Micro-Colonial Fungi (MCFs) which are slow-growing stress-tolerant micro-eukaryotes that specialize in extreme environments. MCFs are paraphyletic and found in the Orders *Chaetothyriales* (*Eurotiomycetes*) and *Dothideales* (*Dothidiomycetes*). We have isolated and described three new MCFs species from desert biological soil crusts (BSCs) collected from two arid land regions: Joshua Tree National Park (Mojave Desert) and UC Natural Reserve at Boyd Deep Canyon (confluence of Mojave and Sonoran Deserts). BSCs are composite assemblages of cyanobacteria, eukaryotic algae, fungi, lichens, and bryophytes embedded into the surface of desert soils, providing a protective buffer against the harsh desert environment. Our work focused on one type of desert BSC, the cyanolichen crust dominated by *Collema sp.* Using culture-dependent protocols, three MCFs were axenically isolated from their respective samples along with the extracted DNA. Their genomes were sequenced using Illumina and Nanopore, and finally assembled and annotated using hybrid assembly approaches and established bioinformatics pipelines to conduct final taxonomic phylogenetic analysis and placement. The three species described here are unique specimen from desert BSCs, here we introduce, *Neophaeococcomyces mojavensis* (*Chaetothyriales)*, *Cladosporium tulheliwenetii* (*Dothideales*), and *Taxawa tesnikishii* (*Dothideales*).

## Introduction

### Biological Soil Crusts

Dryland ecosystems constitute some of the largest terrestrial biomes, collectively covering 47.2% of Earth’s land surface (6.15 billion hectares) (Lal 2004). Drylands are expanding due to climate change and population pressures, while also supporting over 38% of the global human population (Lal 2004; Maestre et al. 2012). Biological Soil Crusts (BSCs) cover up to 70% of dryland ecosystems (Rozenstein and Karnieli 2015). They are an ancient feature of dryland ecosystems, with the earliest known fossil records indicating nearly 2.6 billion years ago (Beraldi-Campesi and Retallack 2016). The initial phases of BSC formation in drylands are the stabilization of the soil surface by filamentous cyanobacteria, followed by the colonization of lichens and bryophytes (Weber et al. 2016). Within dryland ecosystems, the carbon and nitrogen dynamics are often described as the “pulse-reserve model” where precipitation events happen sporadically, stimulating intermittent biological activity which generates biomass and organic matter. After extended periods of heat and dryness, the biological activity goes into reserve and becomes dormant again (Pietrasiak et al. 2013). BSCs also increase soil fertility by increasing soil stability and adding fixed atmospheric carbon and nitrogen to the soil underneath, allowing for greater complex microbial and micro-faunal populations (Housman et al. 2007).

Within BSCs is a consortium of cyanobacteria, algae, diazotrophic bacteria, lichen, fungi, and bryophytes that colonize and stabilize the soil surfaces against natural forces like wind and water (Warren et al. 2019). These microorganisms recruit partners to create a mutualistic symbiotic relationship which helps maintain a self-sustaining micro-ecosystem. Phototrophic organisms fix carbon that is later used by surrounding micro-organisms as an energy source (Green et al. 2016).

The presence of chemoheterotrophic bacteria and cyanobacteria indicates that BSCs are capable of fixing atmospheric nitrogen to increase levels of fixed nitrogen for nearby living vascular plants (Belnap and Harper 1995; Evans and Ehleringer 1993). BSCs have been shown to reduce surface erodibility as filaments of cyanobacterial sheath material lipopolysaccharides (LPS) entangle surface particles and create a crust that is more resistant to entrainment than the layers below (Lal 2004).

Different ecological sites promote different crust taxa. BSCs are classified into types based on the dominant photosynthetic microorganism in the crust which includes: cyanobacteria, algae, lichens, or mosses to produce Cyano-Lichenized Crust, Light Algal Crusts, Dark Algal Crusts, Lichenized Crust, Rough Moss Crust and Smooth Moss Crust (Pietrasiak 2005). The community types are classified based on dominant taxa and their distinct morphologies (Pietrasiak et al. 2013). BSC’s main roles include: 1) mediating almost all inputs and outputs (gasses, nutrients, water) to and from the strata above and below the surface, 2) being the zone of high nutrient deposition, transformation, and availability, 3) structuring temporal, spatial, and compositional aspects of the surrounding vascular plant community, and 4) facilitating the direct delivery of carbon, nutrients, and water by BSCs from the soil interspace to nearby vascular plants.

### Micro-colonial Fungi

‘Black yeasts and relatives’ comprise a large amalgamate of melanized fungi. Among these are micro-colonial fungi (MCF) or rock-inhabiting fungi (RIF), prevalent names used to describe a special group of melanized fungi. Micro-colonial fungi are ubiquitous and found in the most extreme environments. They are present in widely divergent habitats, such as on the indoor rubbers of dishwashers, in oil deposits, on the surfaces of Mediterranean monuments, on the radioactive walls of the Chernobyl Station, in mutilating skin infections in third-world countries, in human superficial and invasive infections, or colonizing lungs of cystic fibrosis patients (de Hoog et al. 2005; Hoog et al. 2000; Vicente et al. 2008; Gümral et al. 2014; Saunte et al. 2012; Thanh and Hien 2019; Mehdiabadi and Schultz; Little and Currie 2008; de Hoog and Hermanides-Nijhof 1977). MCFs are compelling not only because of where we can discover them and their unique physiological features, but also because of how little we know about them.

MCFs have three main characteristics: meristematic growth, presence of 1,8-DHN melanin, and membrane-associated carotenoids and intracellular mycosporine-like amino acids (Staley et al. 1982; Gorbushina et al. 2003). These features distinguish them from other fungi, yet phylogenetically they fall into two paraphyletic Ascomycota clades, the *Dothideomycetes* and *Chaetothyriomycetes*. Melanin is a highly complex secondary metabolite that is produced by many organisms through different evolutionary lineages (Siletti et al. 2017; Wheeler 1983). Most importantly in MCFs, it has been used as a virulence factor as a non-specific armor against host immune system to infect its host (Steenbergen and Casadevall 2003; Casadevall et al. 2003; Casadevall et al. 2000; Nosanchuk et al. 2015; Taborda et al. 2008; Gómez and Nosanchuk 2003). Available research indicates melanin has other functional roles, such as metal chelation, photoprotection, protection from heat and cold stress, binding to antifungals to reduce efficacy, radioprotection (physical shielding and quenching of cytotoxic free radicals), protection from desiccation, interacting with a range of electromagnetic radiation frequencies and mechanical-chemical cellular strength (Dadachova et al. 2008; Casadevall et al. 2003; Gorbushina et al. 2003). Understanding poorly studied fungal function in one extreme environment could yield insight into MCF adaptation in other extreme environments.

The objective of this study was to collect MCF from biological soil crust samples from desert drylands. During our collection, we isolated 3 MCF that had not been described previously. Our efforts to converse with indigenous tribes allowed us to provide a new naming scheme for our strains. We also used a new methodology described herein using BUSCO nucleotide and amino acid sequences to build a multi-locus phylogenetic tree, which proved to be similar to a more conventional tree of MCF which used four highly conserved markers.

## Materials and Methods

### Sampling

Biological soil crusts were collected from two locations in Southern California. The first crust location at the northern edge of Joshua Tree National Park. The second crust location was from Boyd Deep Canyon, near the campgrounds in the UC Natural Reserve. Metadata and coordinates can be found in **Table 1**. All materials were collected under permits from the BLM or permission of the UC National Reserves.

**Table 1.** Three species were collected from biological soil crusts collected from two desert locations in Southern California. Strain JES_112 was isolated from the south east region of Joshua Tree National Park in the Mojave Desert, while JES_115 and JES_119 were both collected from biological soil crusts collected from the campground site in the UC Natural Reserve of Boyd Deep Canyon.

| Species | Date | Location | Coordinates |
| --- | --- | --- | --- |
| JES_112 | November 2017 | Joshua Tree National Park, CA | 34.1026 °N,<br>-115.4542°W |
| JES_115 | July 2019 | Boyd Deep Canyon Reserve, CA | 34.0182 °N,<br>-116.0262 °W |
| JES_119 | July 2019 | Boyd Deep Canyon Reserve, CA | 34.0182 °N,<br>-116.0262 °W |

### Culturing and isolation

Media for isolation, cultivation, and collection used were Malt Extract Yeast Extract (MEYE) media for nutrient-rich complex media and nutrient-poor, oligotrophic media Glucose Asparagine Agar (GAA). The recipe for both media can be found in our protocols.io and the step-by-step protocol for culturing MCFs (Kurbessoian 2019)). The mixing of the sample with the media provides an environment with different layers of dissolved oxygen. Fast-growing fungi require full oxygenation and grow on the surface of the media, while slow-growing MCFs propagate on the bottom of the culture plate, requiring low levels of oxygen. This also allowed a much easier way to culture and select the strains. Once grown, axenic MCFs were isolated by using a flamed loop to pick and place onto clean media.

### Morphological analysis

Strains were grown on Malt Extract Agar (MEA; Oxoid, England), Oatmeal Agar (OA; Oxoid) and Potato dextrose Agar (PDA; Sigma, Germany) incubated at 28°C for 14 days under UV light. Colonies were photographed with a Nikon C-DSD230 dissecting microscope. Microscopic mounts were made in lactic acid and examined using a light microscope (Nikon Eclipse 80i). Pictures were taken with a digital camera (Nikon Digital Sight, DS-5M) and adjusted in Adobe Photoshop CS.

### DNA extraction

Each isolate was grown on MEYE for approximately 1.5 weeks at 22°C. Genomic DNA was extracted from hyphae collected from MEYE plates using the CTAB protocol described in our protocols.io document (Carter-House et al. 2020). Melanin was removed using multiple iterations of phenol:chloroform and chloroform washes of our sample. Genomic DNA was measured by Nanodrop and diluted to ∼28ng/µl. DNA extractions were sent to the UCR Genomics core (Riverside, CA) for 2×150bp sequencing on an Illumina NovoSeq 6000. DNA from all three isolates was also extracted and sequenced on Oxford Nanopore (ONT) platform with library preparation and sequencing following the manufacturer’s directions (Oxford Nanopore, Oxford United Kingdom). Flow cell versions FAK68780, FAQ05493, FAL40904, and FAL35715 were used along with base-calling on the cluster using guppy (*v.3.4.4.+a296cb*) (Wick et al. 2019).

### ITS sequencing

To identify the isolates, extracted DNA was used as a template for PCR amplification using fungal primers ITS1-F (CTTGGTCATTTAGAGGAAGTAA) and ITS4 (TCCTCCGCTTATTGATATGC) along with the PCR setup. PCR was performed in 25 μL reaction volumes using 12.5 μL of Taq 2X MasterMix (NEB), 9.5μL water, 1μL of 20 picomol ITS1-F, 1μL of 20 picomol ITS4 and 1μL of template DNA. Using Bio-Rad thermal cycler, the samples were passed through denaturation and enzyme activation at 95°C for 5 minutes, followed by amplification for 35 cycles at the following conditions: 30 seconds at 95°C, 1 minute at 72°C, and 30 seconds at 65°C. A final 5-minute extension at 72°C completes the protocol. PCR results typically yield between 50-60 ng of amplified DNA.

### Genome assembly and annotation

Initial genome assemblies were constructed for the three MCF isolates using Illumina sequencing. Each MCF was also re-sequenced using Oxford Nanopore technology. All genomes were *de novo* assembled with the AAFTF pipeline (*v.0.2.3*) (Palmer and Stajich 2022) which performs read QC and filtering with BBTools bbduk (*v.38.86)* (Bushnell 2014) followed by SPAdes (*v.3.15.2*) (Bankevich et al. 2012) assembly using default parameters, followed by screening to remove short contigs < 200 bp and contamination using NCBI’s VecScreen. The BUSCO ascomycota_odb10 database (Manni et al. 2021) was used to determine how complete the assembly 3 isolates were. A hybrid assembly of each was generated using MaSUrCa *(v.3.3.4)* (Zimin et al. 2013) as the assembler using both Nanopore and Illumina sequencing reads. General default parameters were used except: CA_PARAMETERS=cgwErrorRate=0.15, NUM_THREADS=16, and JF_SIZE=200000000.

We predicted genes in each near-complete genome assembly with Funannotate (*v1.8.1*) (Palmer and Stajich 2020). A masked genome was created by generating a library of sequence repeats with the RepeatModeler pipeline (Smit and Hubley 2008). These species-specific predicted repeats were combined with fungal repeats in the RepBase (Bao et al. 2015) to identify and mask repetitive regions in the genome assembly with RepeatMasker (*v.4-1-1*) (SMIT 2004). To predict genes, *ab initio* gene predictors SNAP (*v.2013_11_29*) (Korf 2004) and AUGUSTUS (*v.3.3.3*) (Stanke et al. 2006)were used along with additional gene models by GeneMark.HMM-ES (*v.4.62_lic*) (Brůna et al. 2020), and GlimmerHMM (*v.3.0.4*) (Majoros et al. 2004) utilize a self-training procedure to optimize *ab initio* predictions. Additional exon evidence to provide hints to gene predictors was generated by DIAMOND BLASTX alignment of SwissprotDB proteins and polished by Exonerate (*v.2.4.0*) (Slater and Birney 2005). Finally, EvidenceModeler (*v.1.1.1*) (Haas et al. 2008) generated consensus gene models in Funannotate that were constructed using default evidence weights. Non-protein-coding tRNA genes were predicted by tRNAscan-SE (*v.2.0.9*) (Lowe and Chan 2016). The annotated genomes were processed with antiSMASH (*v.5.1.1*) (Blin et al. 2021) to predict secondary metabolite biosynthesis gene clusters. These annotations were also incorporated into the functional annotation using Funannotate. Putative protein functions were assigned to genes based on sequence similarity to InterProScan5 (*v.5.51-85.0*) (Jones et al. 2014), Pfam (*v.35.0*) (Finn et al. 2014), Eggnog (*v.2.1.6-d35afda*) (Huerta-Cepas et al. 2019), dbCAN2 (*v.9.0*) (Zhang et al. 2018) and MEROPS (*v.12.0*) (Rawlings et al. 2018) databases relying on NCBI BLAST (*v.2.9.0+)* (Sofi et al. 2022) and HMMer (*v.3.3.2*) (Potter et al. 2018). Gene Ontology terms were assigned to protein products based on the inferred homology based on these sequence similarity analyses. The final annotation produced by Funannotate was deposited in NCBI as a genome assembly with gene model annotation.

Sequence Read Archive (SRA) files and genome assembly and annotation files can be found under BioProject PRJNA631111.

### Mating type determination

The Mating Type (MAT) locus was identified by searching for homologous MAT genes. The identified homologous regions were examined for their conserved synteny of the MAT locus using clinker (Gilchrist and Chooi 2021) and a custom Biopython script (Cock *et al*. 2009) (Kurbessoian 2022) to extract the annotated region of the genome which contained the locus.

### Telomeric repeat determination

Identification of telomeric repeat sequences was performed using the FindTelomeres.py script (https://github.com/JanaSperschneider/FindTelomeres). Briefly, this searches for chromosomal assembly with a regular expression pattern for telomeric sequences at each scaffold’s 5’ and 3’ end. Telomere repeat sequences were also predicted using the Telomere Identification toolkit (tidk) (*v.0.1.5*) “explore” option (https://github.com/tolkit/telomeric-identifier).

### Inferring phylogenetic relationships of species

Inference of the species relationship was made through phylogenetic trees containing the three new MCF strains described here and available named species with sequence data.

A multi-gene tree was generated using collected BT2 (Beta-Tubulin), CAM (Calmodulin), ITS (Inter-Transcribed Spacer), and LSU (Large Subunit) genes from a total of 32 species. These samples were chosen to compare to the three MCF species with closely related species to confirm identification as a novel lineage. The final selection of 32 species was based on iterations of sequence searching and tree building data analyses to determine close relatives. To include genes from genomes, genome assemblies were searched by TBLASTN tool of NCBI-BLAST (*v.2.9.0+*) using *Zymoseptoria tritici* genes (AAS55060.1 for Beta tubulin) and (AEI69650.1 for Calmodulin). The collection of ITS and LSU genes were obtained from NCBI-Genbank based on BLASTN searches. Each gene file contained FASTA formatted sequences from each species or isolate, the sequences were aligned with MUSCLE (*v.5.1*) and trimmed for low quality alignment regions using Clipkit (*v.1.3.0).* Nucleotide substitution models were determined for each alignment with IQTREE2. The alignments were combined into a single ‘supermatrix’ alignment with combine_multiseq_aln.py script provided in PHYling. The script also produced a partition file listing the sub-locations of each marker gene alignment in the super-alignment. IQTree2 was run to find a substitution model fit for and partition merging. To reduce the computational burden that occurs with including a large number of organism ribosomal sequences we used the relaxed hierarchical clustering algorithm (Lanfear et al. 2014) using the tag -m MF+MERGE and running 1000 bootstrapping on the alignment.

### Choosing to work with indigenous tribes

Our choice for naming of the three MCFs isolated from BSCs is rooted in the traditional indigenous land where these were found. *Neophaeococcomyces mojaviensis* was isolated from BSCs collected from the south-east region of Joshua Tree National Park, territories ancestral to Yuhaaviatam/Maarenga’yam (Serrano) and Newe Segobia (Western Shoshone) Tribes (Brown and Boyd 1922; Kroeber 1925). The isolates of *Coniosporium tulheliwenetii* and *Taxawa tesnikishii* were isolated from BSCs collected from UC Reserve Boyd Deep Canyon (Reserve doi:10.21973/N3V66D), land of the traditional ancestral Cahuilla Tribe (White 2001; Hooper 1920). The word *tulheliweneti* in Cahuilla means, black-spread-be-noun, while *Taxaw* means body, and *tesnikish* means yellow. In Cahuilla traditions, Taxaw tesnikish is the deity that resided in Boyd Deep valley. To recognize the origins, we consulted with Tribal linguists and speakers to derive their names from Cahuilla and Serrano terms that describe either geographic region, oral tradition, or features that help distinguish them.

## Results

### Biological soil crust collection

Biological soil crusts were collected from the two regions described in **Table 1**. The method of sterile collection is through the use of sterile empty Petri dishes, stamped on top, with a sterile spatula used to grab the soil from underneath, flip the sample into the dish, place the lid on top and seal it with parafilm.

### Culturing results

Samples material was used to culture fungi using a collection of techniques in our laboratory at UC Riverside. More than 50+ fungal culture specimens were collected from biological soil crusts, 18 were MCF cultures. The pour plate protocol used allows for fast-growing fungi to appear on the top, while MCF grows within the bottom layer of media in Petri dishes. Samples were excised from deep in the media and allowed to grow on fresh MEYE. Each sample had the DNA extracted and each strain ITS was Sanger sequenced to determine the identity. JES_112 was initially identified as a *Knufia* species. JES_115 and JES_119 were identified as *Dothideales* species, but there remained ambiguity on the exact species identification as a near perfect sequence match was not found for ITS1 to assign a name.

### Sequencing and assembly of MCF isolates

To gain information on the genome content for the recovered MCF isolates, we sequenced and assembled the genomes of the three isolates (**Table 2**). Each isolate was sequenced with Illumina MiSeq and Oxford Nanopore (ONT) technology and constructed into a hybrid assembly that combined both read types (**Table 2**). The depth of coverage ranged from 25-43x Illumina coverage across the three specimens, while Nanopore ranged from 0.83-194.58x. Assessment of the genome completeness was performed with BUSCO (Benchmarking Universal Single-Copy Orthologs) and results indicated the genomes had ranges of 73-92.9% complete gene copies found, with 73-92.8% of the markers appearing as single copies in the assembly, a low fraction of duplication (0.1-0.2). Contig counts ranged from 38-480, while the average genome assembly size is about 30 Mbps for all isolates. The L50 statistic, a measure of the count of the smallest number of contigs whose sum makes up half of the genome size, ranged from 9-88, and the N50 statistic, a measure of the smallest contig length for covering half of the genome size, ranging from 117kbp −1.232 Mbp. To further assess completeness, we tested for the presence of telomeric repeat units “TTTAGGG/CCCTAA”. These were identified as repeat arrays at both ends of contigs in JES_115 and JES_119, 61 total telomeres and 55 total telomeres respectively, though the results were inconclusive for JES_112. No telomeric repeat-containing contigs were detected in JES_112 which could be due to the fragmented nature of the genome assembly. The fragmented assembly also is reflected in the lower BUSCO score for this genome (**Table 2**).

**Table 2.** Genome assembly and annotation summary statistics. All strains were sequenced with Illumina and Nanopore sequencing. With the added sequencing reads, we were able to glean telomere information along with genome completeness. It is clear that JES_112 is contaminated as the BUSCO score is lower, while JES_115 and JES_119 are much closer to a more complete genome.

|  | **JES_112** | **JES_115** | **JES_119** |
| --- | --- | --- | --- |
| Scaffold Count | 478 | 64 | 38 |
| Total Length | 32,593,754 bp | 32,607,911 bp | 29,383,553 bp |
| Minimum Length | 561 | 4823 | 1783 |
| Maximum Length | 427,480 | 1,739,523 | 2,828,030 |
| Mean Scaffold Length | 68,187.77 | 509,498.61 | 773,251.39 |
| Scaffold L50 | 87 | 14 | 9 |
| Scaffold N50 | 117,971 | 966,560 | 1,232,081 |
| Scaffold L90 | 282 | 31 | 24 |
| Scaffold N90 | 32,261 | 507,175 | 558,687 |
| BUSCO Complete %<br>Illumina | 84.7 | 90.2 | 97.4 |
| BUSCO Complete %<br>Nanopore | 73.4 | 92.6 | 92.9 |
| BUSCO Duplicate %<br>Nanopore | 0.2 | 0.1 | 0.1 |
| BUSCO Single %<br>Nanopore | 73.2 | 92.5 | 92.8 |
| GC% | 48.25 | 52.55 | 51.98 |
| Illumina coverage | 34.58x | 43.45x | 25.38x |
| Nanopore coverage | 0.83x | 20.44x | 194.58x |
| Telomere Count | 0, 0F, 0R | 61, 29 F, 32 R | 55, 29 F, 26 R |
| Number of Genes | 10,996 | 9,304 | 9,386 |
| mRNA genes | 10,924 | 9,226 | 9,189 |
| tRNA genes | 72 | 78 | 197 |
| GO Terms | 6,001 | 5,089 | 5,321 |
| InterProScan | 8,523 | 7,079 | 7,291 |
| EGGNOG | 9,934 | 8,487 | 8,511 |
| PFAM | 7,520 | 5,982 | 6,219 |
| CAZYme | 309 | 257 | 346 |
| MEROPS | 344 | 284 | 291 |
| Secretion | 674 | 532 | 631 |

The number of predicted tRNA genes ranged from 76-197, the total number of protein-coding genes predicted ranged from 9,029-10,920, and secondary metabolite cluster counts ranged from 15-27, 27 resulting from JES_112.

### Mating type loci determination

The mating types were determined using prior work on members of Chaetothyriales and Dothideomycetes including *Exophiala dermatitidis*, and *Aureobasidium pullulans* (Metin et al. 2019; Teixeira et al. 2017). Using collected nucleotide and protein FASTA sequences, BLAST was used to identify the genes found in these loci. The genes in this locus are generally flanked by SLA2 and APN2 and may sometimes have alternative genes within. **Figure 2** was generated using JES_112, *Exophiala aquamarina* CBS 119918, *E. dermatitidis* NIH/UT8656, *E. dermatitidis* CBS 115663, and *Capronia coronata* CBS 617.96. The MAT locus is flanked by two genes, SLA2 (purple) and APN2 (orange). The MAT 1-1 gene, MAT 1-1-4 (green), and MAT 1-1-1 (teal) are observed in *E. dermatitidis* CBS 115663 and *Capronia coronata*, while *E. dermatitidis* NIH/UT8656 contains MAT 1-2 (pink) as does *Capronia coronata*. Our organism in question, *Neophaeococcomyces mojaviensis* contains the MAT 1-2 gene along with the MAT 1-1-1 gene. This is similarly seen with *N. aloes* FJII-L3-CM-P2. This is confusing as the MAT 1-1-1 gene usually pairs with MAT 1-1-4. The gene arrow in blue is indicated as an L-type calcium channel domain, while the gene in yellow is an unidentified functional gene or domain.

**Figure 1.**
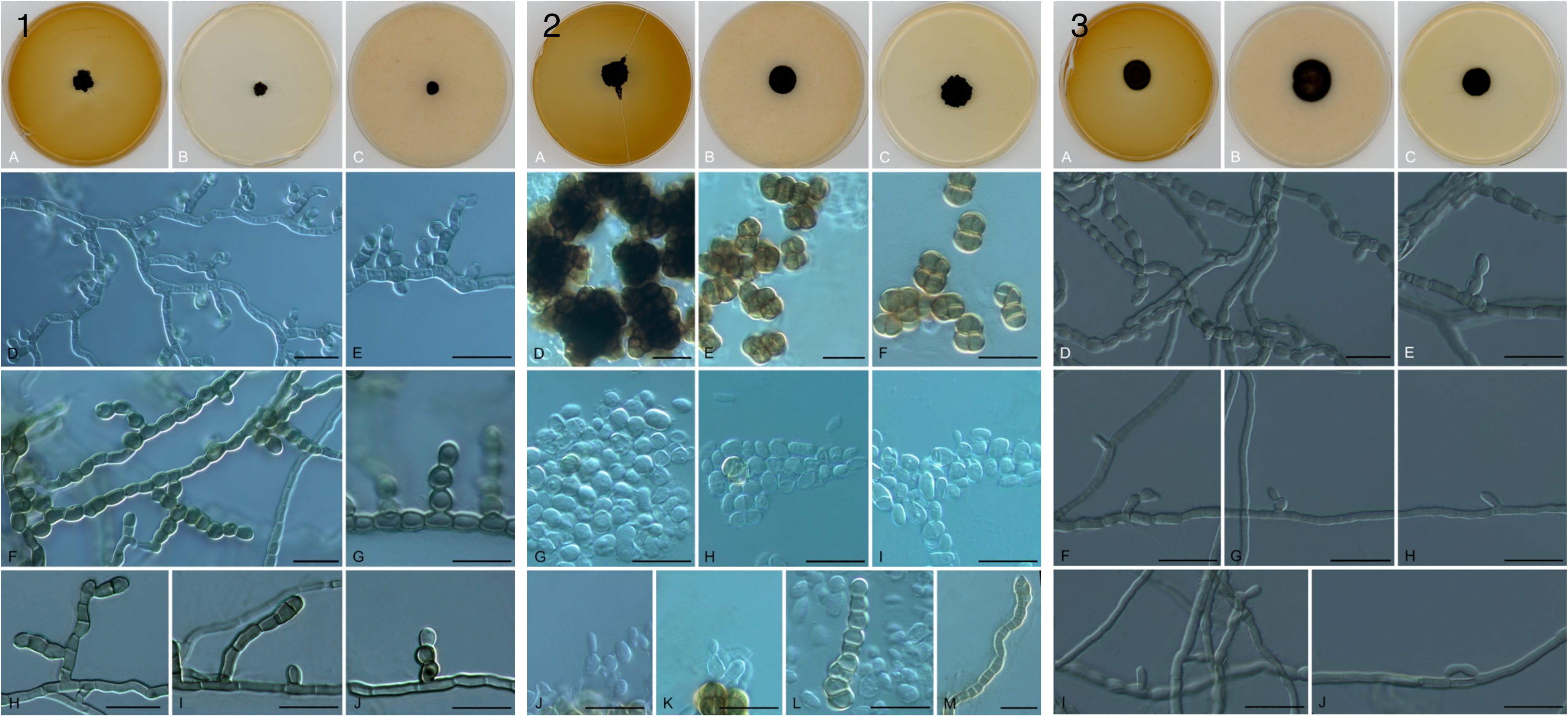
Microscopic morphology of isolated MCF. (1) *Taxawa tesnikishii* JES_119, colonies on MEA (A), OA (B), and PDA (C) agars, meristematic cells (D-F), and budding cells (G-M); (2) *Coniosporium tulheliwenetii* JES_115, colonies on MEA (A), OA (B), and PDA (C) agars, hyphae with lateral branches of swollen cells reluctantly leading to arthroconidia (D-J); (3) *Neophaeococcomyces mojaviensis* JES_112, colonies on MEA (A), OA (B), and PDA (C) agars, hyphae producing intercalary swollen cells without conidiation (D-J). Microscopic mounts were made in lactic acid and examined using a light microscope. The scale depicted here is 25µm.

**Figure 2.**
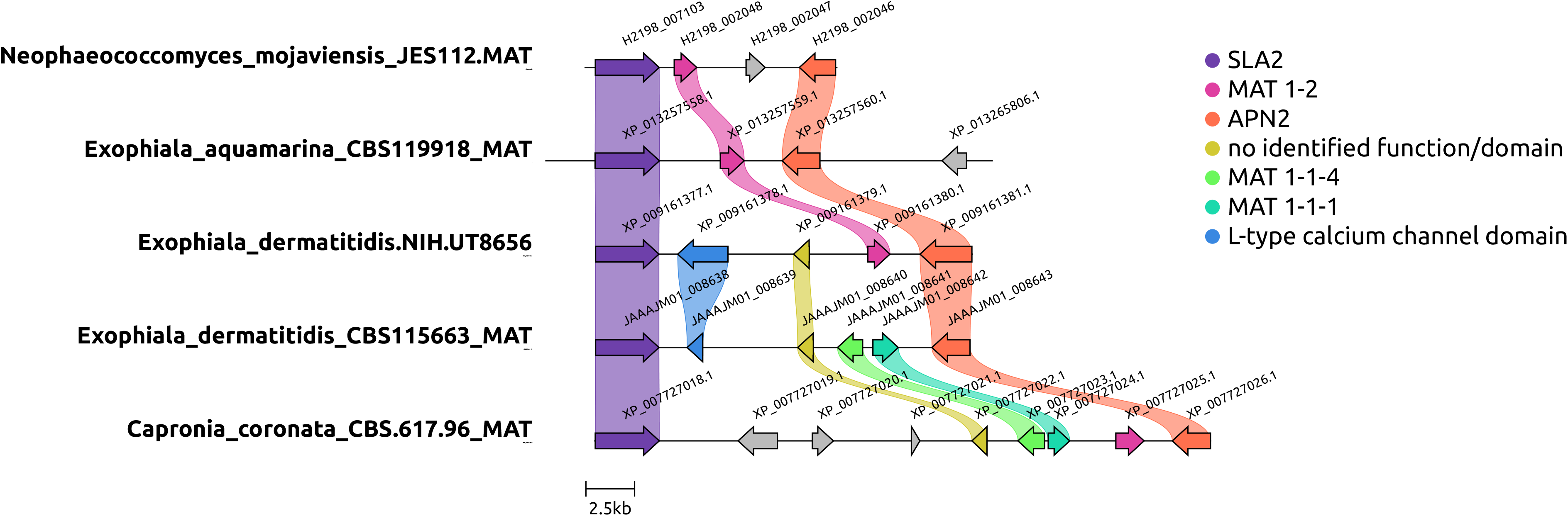
Mating type loci determined for Chaetothyriales JES_112 here called *Neophaeococcomyces mojaviensis*. Other MCFs added to this analysis include *Neophaeococcomyces aloes* FJII-L3-CM-P2*, Exophiala dermatitidis* NIH/UT8656, *E. dermatitidis* CBS 115663, and *Capronia coronata* CBS 617.96. The MAT locus is flanked by two genes, SLA2 (purple) and APN2 (orange). The MAT 1-1 gene, MAT 1-1-4 (green), and MAT 1-1-1 (teal) are observed in *E. dermatitidis* CBS 115663 and *C. coronata*, while *N. aloes* and *E. dermatitidis* NIH/UT8656 contain MAT 1-2 (pink) as does *C. coronata*. Our organism in question, *N. mojaviensis* contains the MAT 1-2 and MAT 1-1-1 gene which is confusing as it is not common to lose the MAT 1-1-4. The gene arrow in blue is indicated as an L-type calcium channel domain, while the gene in yellow is an unidentified functional gene or domain.

Figure 3 was generated using JES_115 and JES_119 and two other species including, *Aureobasidium pullulans* EXF-150 and *Coniosporium apollinis* CBS 100218. All species described here only have the MAT 1-2 gene. The MAT locus is flanked by two genes, SLA2 (purple) and APN2 (orange). *Dothideomycetes* mating type loci look very different from the *Chaetothyriales* mating type loci. Other genes included in this locus concern only a single MAT locus, MAT 1-2 (pink), and rRNA adenine demethylase (yellow). Certain genes like the AP endonuclease (teal) and a hypothetical protein (blue) are not seen in all four genomes.

**Figure 3.**
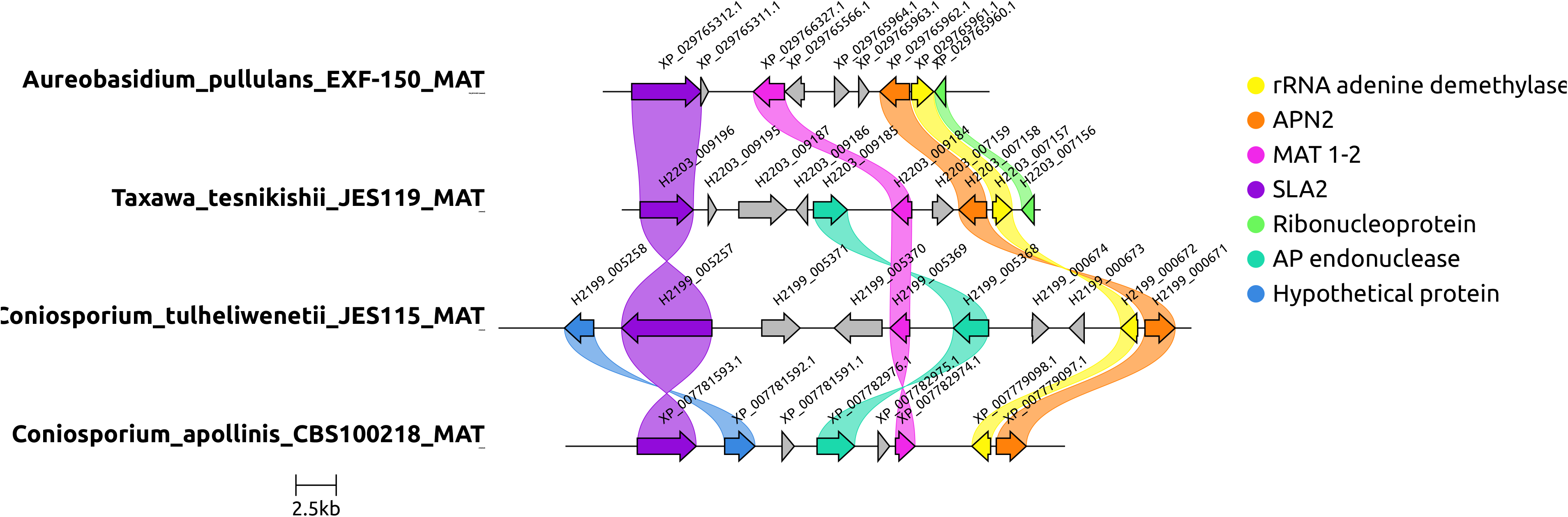
Mating type loci determined for Dothideomycetes JES_115 and JES_119, here called *Coniosporium tulheliwenetii* and *Taxawa tesnikishi*. Other dothideaceous black yeast-like fungi in this analysis include *Aureobasidium pullulans* EXF-150, and *Coniosporium apollinis CBS* 100218. All species described here only have the MAT 1-2 gene. The MAT locus is flanked by two genes, SLA2 (purple) and APN2 (orange). *Dothideomycetes* mating type loci look very different from the *Chaetothyriales* mating type loci. Other genes included in this locus include only a single MAT locus, MAT 1-2 (pink). Another gene included in this locus is rRNA adenine demethylase (yellow). Certain genes like the AP endonuclease (teal) and a hypothetical protein (blue) are not seen in all four genomes.

### MCF morphology

Strains showed limited and slow growth with black colonies on all media tested. Strain JES_119 produced abundant unilateral, hyaline budding cells which soon inflated, developed cruciate septation, and became melanized. Strain JES_115 was hyphal with lateral chains of swollen cells which occasionally detached. Strain JES_112 lacked sporulation, the thallus consisting of melanized densely septate hyphae with series of somewhat inflated cells

During the annotation step, we processed the genomes through antiSMASH (**Table 3**), to detect Biosynthetic Gene Clusters (BGCs) of secondary metabolites from the annotated genome sequence. Once predicted metabolite of note, beta-lactone was found in all three MCF. A further literature search indicates the importance of beta-lactone in industry as there have been multiple products derived from these chemicals including, antifungals and anti-cancer agents (Robinson et al. 2019). A BLASTP search of the JES_112 beta-lactone gene identified homologs to Acetyl-CoA synthetase + 2-isopropylmalate synthases + bile acid dehydrogenase, zinc-binding alcohol dehydrogenase + acetoacetate-CoA + NAD-dependent alcohol dehydrogenase, and AMP-binding protein + 2-isopropylmalate synthases.

**Table 3.** Predicted secondary metabolite gene clusters. The BGCs predicted include multiple copies of PKS and NRPS genes, but siderophores were only recovered in JES_112. There was a higher number of BGCs predicted in JES_112 (27) our *Chaetothyriales* species, than JES_115 and JES_119 (15) our *Dothideomycetes*. This table lists the numbers of each. JES_112 has a much higher number (27) of secondary metabolites than the other two (15).

|  | T1PKS | Terpene | T3PKS | NRPS | NRPS-like | Siderophore | Arylpolyene | Betalactone | Fungal RiPP |
| --- | --- | --- | --- | --- | --- | --- | --- | --- | --- |
| JES 112 | 6 | 6 | 2 | 4 | 6 | 1 | 1 | 1 | 0 |
| JES 115 | 5 | 4 | 0 | 2 | 3 | 0 | 0 | 1 | 0 |
| JES 119 | 1 | 4 | 0 | 1 | 7 | 0 | 0 | 1 | 1 |

**Table 4.** Gene Tree Nucleotide Accession numbers. Collected accession numbers of alternative strains to build Figures 4 and 5. The remaining empty spaces indicate the need to pull the sequences using BLAST. Once collected each gene populated a gene FASTA file, aligned, clipped and IQTREE2, a tree-building program, was run to determine the gene models for each gene. Once determined, the gene alignment lengths were calculated and partitioning scripts were described and used to build a larger tree, using IQTREE2.

| Species | Beta-Tubulin | Calmodulin | Internal-Transcribed Spacer | Large Subunit (Ribosomal) |
| --- | --- | --- | --- | --- |
| *Neophaeococcomyces mojaviensis* JES112 | - | - | - | - |
| *Taxawa tesnikishii* JES119 | - | - | - | - |
| *Alternaria solani* NL03003 | MK388240.1 | MH243808.1 | NR 172408.1 | ON416546.1 |
| *Anthraccina ramosa* CGMCC3.16372 | KP226556.1 | - | NR 172152.1 | KP174923.1 |
| *Anthraccina saxicola* CGMCC3.17315 | KP226554.1 | - | NR 172151.1 | KP174921.1 |
| *Arthrocladium caudatum* CBS457.67 | LT558710.1 | - | NR 145398.1 | MH870732.1 |
| *Arthrocladium tardum* CBS 127021 | - | - | NR 154706.1 | NG 057089.1 |
| *Arthrocladium tropicale* CBS 134926 | KX822406.1 | - | NR 154724.1 | NG 057119.1 |
| *Aureobasidium mangrovei* IBRCM30265 | XM 0299048 52.1 | XM 04102580 9.1 | NR 174637.1 | NG 078639.1 |
| *Aureobasidium* sp.AWRI4619-CO-2020 | XM 0299048 52.1 | XM 04102580 9.1 | MK256457.1 | MK968773.1 |
| *Baudoinia panamericana* UAMH10762 | XM 0076738 85.1 | XM 00767526 4.1 | NR 153615.1 | KT186497.1 |
| *Bradomyces alpinus* CCFEE 5493 | LN589970.1 | - | NR 132844.1 | NG 058641.1 |
| *Bradomyces yunnanensis*<br>CGMCC 3.17350 | KP226545.1 | - | KP174865.1 | KP174926.1 |
| *Capronia coronata* CBS<br>617.96 | XM 0077289<br>82.1 | XM 00772155<br>6.1 | NR 154745.1 | NG 067831.1 |
| *Cenococcum geophilum*<br>1.58 | - | - | KC967410.1 | MN817694.1 |
| *Cladophialophora*<br>*carrionii* CBS 260.83 | KF928582.1 | XM 00872400<br>0.1 | MH861582.1 | MH873312.1 |
| *Cladophialophora yegre*<br>*sii* CBS 114405 | EU137209.1 | - | NR 111284.1 | KX822323.1 |
| *Cladophialophora*<br>*immunda* CBS 834.96 | EU137203.1 | - | NR 111283.1 | MH874242.1 |
| *Coniosporium apollinis*<br>CBS 100218 | - | XM 00778549<br>5.1 | NR 159787.1 | GU250898.1 |
| *Coniosporium uncinatum*<br>CBS 100219 | - | - | NR 145343.1 | NG 058806.1 |
| *Coniosporium uncinatum*<br>CCFEE 6149 | - | - | MZ573424.1 | MZ573424.1 |
| *Cyphellophora aestiva*<br>CBS 239.91 | MW297547.1 | - | MH861291.1 | MH873056.1 |
| *Cyphellophora laciniata*<br>CBS 239.91 | KF928604.1 | - | NR 121335.1 | MH873933.1 |
| *Cyphellophora sessilis*<br>KNU-002 | KX431943.1 | - | KX431941.1 | KX431942.1 |
| *Cyphellophora*<br>*vermispora* CBS 228.86 | KC455227.1 | - | KC455244.1 | KC455257.1 |
| *Exophiala dermatitidis*<br>NIH.UT8656 | - | XM 00916045<br>0.1 | JH226131.1 | ON543213.1 |
| *Exophiala oligosperma*<br>CBS 265.49 | EF551507.1 | XM 01640456<br>0.1 | MH856519.1 | MH868049.1 |
| *Exophiala oligosperma*<br>CBS 127587 | <u>KF928570.1</u> | - | <u>MH864631.1</u> | <u>MH876068.1</u> |
| *Exophiala radialis* P2854 | <u>KT723463.1</u> | - | <u>NR 158397.1</u> | <u>NG 069319.1</u> |
| *Exophiala xenobiotica*<br>CBS 118157 | <u>NW 0135624</u><br><u>90.1</u> | <u>XM 01345465</u><br><u>6.1</u> | <u>OP132917.1</u> | <u>MH875251.1</u> |
| *Fonsecaea pedrosoi* CBS<br>271.37 | <u>XM 0134308</u><br><u>09.1</u> | <u>XM 01342686</u><br><u>4.1</u> | <u>MT023597.1</u> | <u>KF155185.1</u> |
| *Friedmanniomyces*<br>*endolithicus* CCFEE<br>5311 | - | - | <u>NR 144917.1</u> | <u>GU250367.1</u> |
| *Friedmanniomyces*<br>*simplex* CCFEE 5184 | - | - | <u>NR 155079.1</u> | <u>GU250368.1</u> |
| *Hortaea werneckii* EXF-2000 | - | - | <u>NR 145338.1</u> | <u>NG 057773.1</u> |
| *Knufia petricola*<br>MA5789 | - | - | <u>NR 111871.1</u> | <u>MT126669.1</u> |
| *Knufia mediterranea*<br>CCFEE 6211 | - | - | <u>KP791793.1</u> | <u>KR781080.1</u> |
| *Knufia mediterranea*<br>CCFEE 6205 | - | - | <u>KP791794.1</u> | <u>KR781081.1</u> |
| *Knufia separata*<br>CGMCC 3.17337 | <u>KP226564.1</u> | - | <u>NR 172233.1</u> | <u>KP174931.1</u> |
| *Knufia tsuneda*<br>FMR10621 | - | - | <u>NR 132842.1</u> | <u>HG003672.1</u> |
| *Knufia walvisbayicola*<br>CBS 146989 | <u>MZ078266.1</u> | - | <u>NR 173046.1</u> | <u>NG 076737.1</u> |
| *Knufia* sp. MV-2018<br>MLT-8 | - | - | <u>MT636977.1</u> | <u>MT636937.1</u> |
| *Leptosphaeria biglobosa*<br>CA1 | <u>AY749005.1</u> | - | <u>NR 111620.1</u> | <u>JX681093.1</u> |
| *Leptosphaeria maculans*<br>JN3 | <u>AY749012.1</u> | <u>XM 00384071</u><br><u>9.1</u> | <u>M96384.1</u> | <u>JF740306.1</u> |
| *Lithohypha aloicola*<br>CPC35996 | MN556837.1 | - | NR 166313.1 | NG 068316.1 |
| *Lithohypha guttulata*<br>FMR 20100 | OX346320.1 | - | OX346376.1 | OX346377.1 |
| *Metulocladosporiella musicola* CBS 110960 | - | MG934537.1 | MH862870.1 | NG 067447.1 |
| *Metulocladosporiella samutensis* CBS 143921 | - | MG934540.1 | NR 161056.1 | MG934453.1 |
| *Neohortaea acidophila*<br>CBS 113389 | XM 0337332<br>87.1 | XM 03373393<br>0.1 | NR 169648.1 | OL739260.1 |
| *Neophaeococcomyces aloes* FJII-L3-CM-P2 | - | - | NR 132069.1 | KF777234.1 |
| *Neophaeococcomyces catenatus* CBS 650.76 | - | - | AF050277.1 | MH872793.1 |
| *Neophaeococcomyces oklahomaensis*<br>EMSL3313 | - | - | MW798187.1 | MW798182.1 |
| *Phaeosphaeria* sp.MPI-PUGE-AT-0046c | - | - | NR 155675.1 | NG 068582.1 |
| *Polychaeton citri* CBS<br>116435 | - | - | GU214649.1 | OM238161.1 |
| *Polyplosphaeria fusca*<br>CBS 125425 | - | - | NR 119397.1 | MH875147.1 |
| *Salinomyces thailandica*<br>CCFEE 6315 | - | - | NR 171710.1 | MH875167.1 |
| *Scorias spongiosa*<br>HLSS1 | - | - | NR 154422.1 | MH866910.1 |
| *Strelitziana africana*<br>ICMP 21758 | - | - | MK210538.1 | MK210499.1 |
| *Strelitziana africana*<br>ICMP 21760 | - | - | MK210540.1 | MK210501.1 |
| *Teratosphaeria gauchensis* CMW 17545 | - | <u>KF902726.1</u> | <u>NR 144976.1</u> | <u>EU019295.2</u> |
| *Trichomerium cicatricatum* CGMCC3.17307 | <u>KP226540.1</u> | - | <u>NR 172232.1</u> | <u>KP174947.1</u> |
| *Trichomerium deniquilatum* MFLUCC10-0884 | - | - | <u>NR 132965.1</u> | <u>NG 059479.1</u> |
| *Trichomerium gleosproum* MFLUCC10-0087 | - | - | <u>JX313656.1</u> | <u>JX313662.1</u> |
| *Trichomerium* sp. XDY-2021a-IFRDCC3302 | - | - | <u>MN901488.1</u> | <u>MZ822131.1</u> |
| *Tuber melanosporum* | <u>MZ458420.1</u> | <u>XM 00284118 4.1</u> | <u>GQ917052.1</u> | <u>JF807486.1</u> |
| *Venturia pyrina* ICMP11032 | <u>EU853839.1</u> | - | <u>NR 169905.1</u> | <u>AY849967.1</u> |
| *Zalaria obscura* FKI-L7-BK-DRAB2 | - | - | <u>NR 153466.1</u> | <u>MT103097.1</u> |
| *Zymoseptoria tritici* CBS 100329 | <u>XM 0038567 27.1</u> | <u>KF254299.1</u> | <u>KP852517.1</u> | <u>MK788267.1</u> |

A second predicted metabolite of note was the siderophore gene cluster. These genes are involved in iron transport, is a more well-known metabolite that could provide the organism with virulence and protection by sequestering iron molecules from its surrounding environment (Johnson 2008; Sass et al. 2019) and was primarily detected in JES_112.

Additionally, the BSC search revealed fungal-RiPPs (Ribosomal and post-transcriptionally modified proteins) in JES_119. The homolog was best classified as a gamma-glutamyltransferase + cytochrome protein. Fungal-RiPPs have been used as an alternative source of bioactive metabolites and could be of industrial interest (Vogt and Künzler 2019). Additional BGCs were found including Type-1 Polyketide synthases (T1PKS), Type-3 Polyketide synthases, non-ribosomal peptide synthases, non-ribosomal peptide synthase-like, and terpenes.

### Four-locus gene trees

The multi-gene internal-transcribed space (ITS1), large ribosomal subunit (LSU) beta-tubulin, calmodulin tree (Figure 4 and Figure 5) places the new isolates in context of named species in the known genera (**Table 3**).

**Figure 4.**
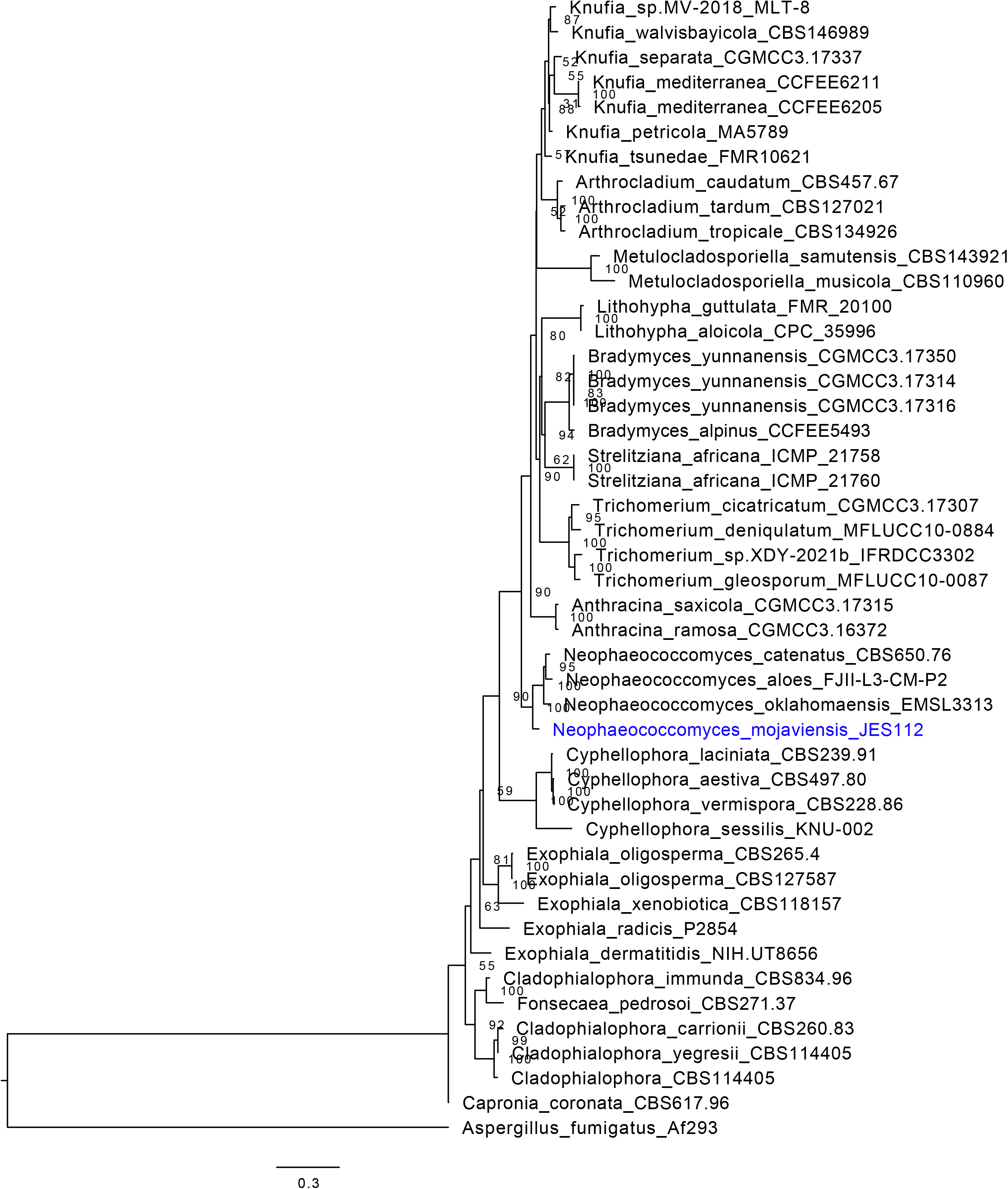
Chaetothyriales multi-locus gene tree. Multi-loci phylogenetic tree using 4 highly conserved genes including, beta-tubulin, calmodulin, internal-transcribed sequence (ITS), and the large subunit of the Ribosome (LSU). Seven representative strains with loci were used, which were aligned, trimmed, and concatenated to run the IQTree program and generate a concise multi-loci gene tree for *Neophaeococcomyces mojaviensis* JES112. Here this strain is placed in the *Neophaeococcomyces* genera.

**Figure 5.**
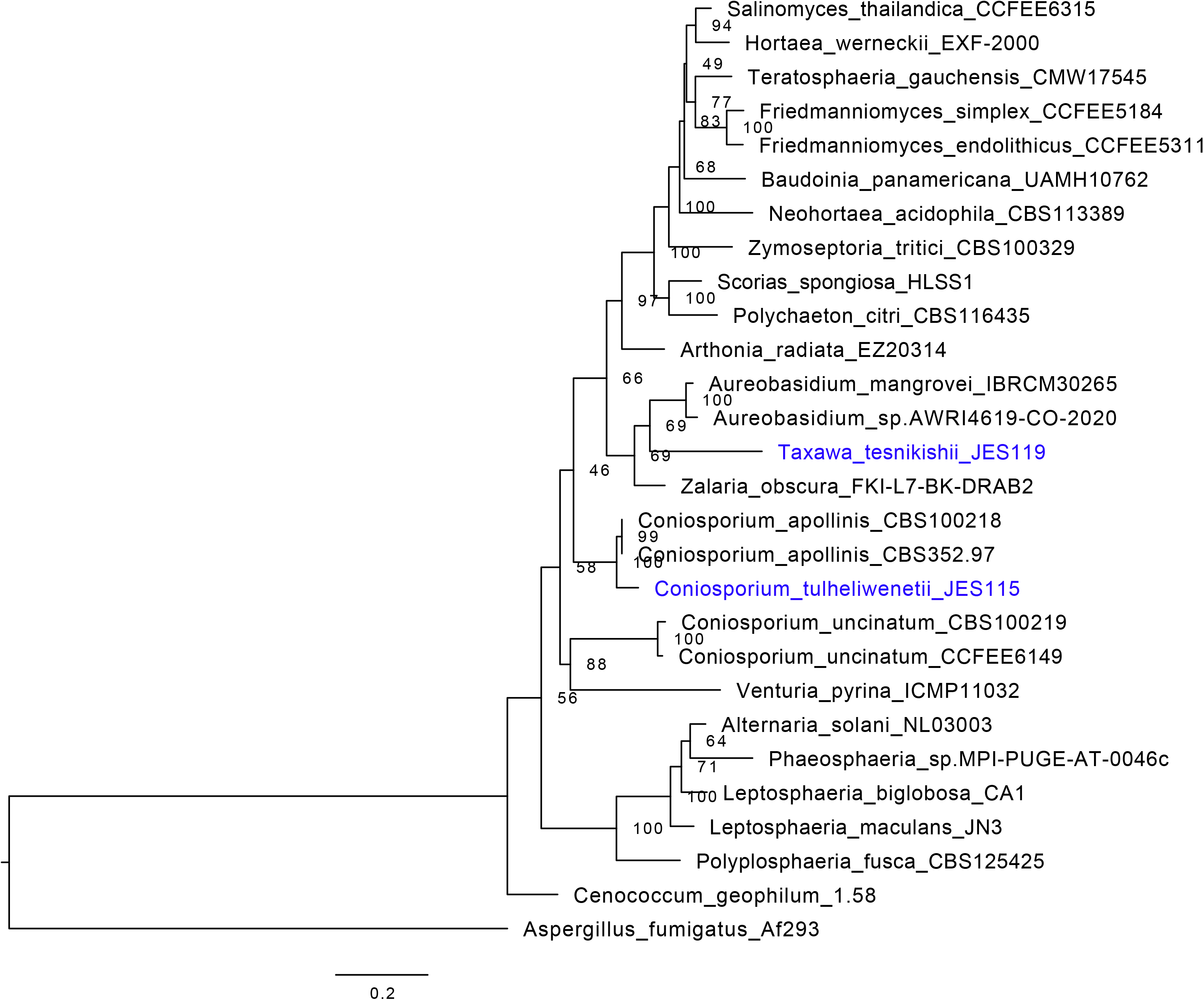
Dothideomycetes multi-locus gene tree. Multi-loci phylogenetic tree using 4 highly conserved genes including, beta-tubulin, calmodulin, internal-transcribed sequence (ITS), and the large subunit of the Ribosome (LSU). Twenty-five representative strains with loci were used, which were aligned, trimmed, and concatenated to run the IQTree program and generate a concise multi-loci gene tree for JES_115 and JES_119 or *Coniosporium tulheliwenetii* and *Taxawa tesnikishii* respectively. Here strain JES_115 is grouping with other *Coniosporium* spp., while JES_119 is grouping with but distinctly apart from *Aureobasidium* spp.

Forty-three isolates were used, and the alignment of the sequences were trimmed, and concatenated to run the IQTree program and generate a concise multi-loci gene tree for *Neophaeococcomyces mojaviensis* JES112. Here this strain is placed in the genus *Neophaeococcomyces*, along with *Exophiala dermatitidis* NIH/UT8656, *E. radicis* P2854, *E. oligospoerma* CBS 265.49, *E. oligospoerma* CBS 127587, and *E. xenobiotica* CBS 11817, *Fonsecaea pedrosoi* CBS 271.37, *Capronia coronata* CBS 617.96, *Cladophialophora immunda* CBS 834.96, *C. carrionii* CBS 260.83, *C. yegresii* CBS 114405, *Neophaeococcomyces aloes* FIJI-L3-CM-P2*, N. catenatus* CBS650.76*, N. oklahomaensis* EMSL 3313*, Knufia petricola* MA5789*, K. separata* CGMCC3.17337*, K. mediterranea* CCFEE 6211*, K. mediterranea* CCFEE6205*, Knufia* sp. MV-2018-MLT-8*, K. walvisbayicola* CBS 146989*, K. tsunedae* FMR10621*, Bradymyces yunnanensis* CGmCC3.17350*, B. alpinus* CCFEE 5493*, Strelitziana africana* ICMP 21758 and ICMP 21760, and *Tuber melanosporum* Mel8 as the outgroup.

The third phylogenetic tree, Figure 5, uses the same method described above as in Figure 4. 25 strains of Dothideomycetes MCF were included to place JES_115 and JES_119 into their proper clades. MCF strains included a collection of *Coniosporium apollinis* CBS 100218, CBS 352.97*, Aureobasidium mangrovei* IBRCM30265*, Aureobasidium* sp. AWRI4619-CO-2020*, Zalaria obscura* FKI-L7-BK-DRAB2*, Polyplosphaeria fusca* CBS 125425*, Leptosphaeria maculans* JN3, *Leptosphaeria biglobosa* CA1*, Phaeosphaeria* sp. MPI-PUGE-AT-0046c*, Alternaria solani* NL03003*, Cenococcum geophilum* 1.58*, Venturia pyrina* ICMP11032*, Coniosporium uncinatum* CBS 100219, and CCFEE 6149, *Scorias spongiosa* HLSS1*, Polychaeton citri* CBS 116435*, Zymoseptoria tritici* CBS 100329*, Neohortaea acidophila* CBS 113389*, Salinomyces thailandica* CCFEE 6315*, Hortaea werneckii* EXF-2000*, Baudoinia panamericana* UAMH 10762*, Teratosphaeria guachensis* CMW 17545*, Friedmanniomyces simplex* CCFEE 5184*, Friedmanniomyces endolithicus* CCFEE 5311, and *Tuber melanosporum* Mel8 as the outgroup. JES_115 groups near *Coniosporium* spp.*, w*hile JES_119 groups near *Aureobasidium* spp.

### Taxonomy

#### *Neophaococcomyces mojaviensis sp. nov.*, Kurbessoian, Ahmed, de Hoog and Stajich -Mycobank (848161). Description

Colonies on MEA, OA, PDA after two weeks of incubation at 25°C attain diameters of 1.6, 2, 2.2, and 1.6 mm respectively. Hyphae 2.9-5.4 μm diameter, dark brown or black, smooth or occasionally rough walled, cells eventually inflating up to 8 μm in diameter. The proliferation of inflated cells might result in the formation of multicellular bodies terminal or intercalary in the hyphae measuring 13.8-31.8 μm, multiseptate with transverse and longitudinal septa; bodies round or irregular, dark brown to black and thick-walled. No conidia were observed. Sexual state unknown.

#### Holotype

ATCC (TSD-444), preserved at the herbarium collection of the Westerdijk Fungal Biodiversity Institute, Utrecht, The Netherlands. Living culture ex-type ATCC (TSD-444).

#### Diagnosis

Strain JES_112 clusters in the order *Chaetothyriales*, relatively close to the numerous rock-inhabiting species of the genus *Knufia*, and amidst the species of the small genus *Neophaeococcomyces*. Both genera are assigned to the family *Trichomeriaceae*, containing ‘sooty molds’ with oligotrophic lifestyles colonizing inert surfaces (Chomnunti et al. 2012). The cluster *Neophaeococcomyces,* based on *N. catenatus* as generic type species, as yet contains only four species, which mostly were isolated from low nutrient habitats, such as the bark of *Aloe*, stone, or air. Several species of the genus show budding with melanized cells, and frequently cells cohere in short chains. Often lateral branches of short inflated cells are inserted on hyphae; the cells may be released as arthroconidia which then swell or appear yeast-like, or remain coherent.

### *Coniosporium tulheliwenetii sp. nov*., Kurbessoian, Ahmed, de Hoog and Stajich -MycoBank (848162)

#### Description

Colonies on MEA, OA, PDA, after 2 weeks of incubation at 25°C attain diameter restricted, measuring 1.1 cm, 0.72 cm, and 0.75cm, respectively. Hyphae 4.6-8.4 μm diameter showing lateral chains of swollen cells 7.7 - 10.8 μm. The cells however remain intact, and no conidia was observed. Teleomorph unknown.

#### Holotype

**CBS (150075), and herbarium CBS H-25288**, preserved at the herbarium collection of the Westerdijk Fungal Biodiversity Institute, Utrecht, The Netherlands. Living culture ex-type CBS (150075).

#### Diagnosis

Strain JES_115 was morphologically similar to *Knufia* species in the presence of lateral chains of swollen cells inserted on main hyphae, the cells eventually being released as conidia, but was phylogenetically remote. It clustered with some species of rock-inhabiting described in *Coniosporium,* in the order *Pleosporales*. Interestingly, the original descriptions, which were made before sequence data were available, classified them all into a single genus. The genus *Coniosporium* is one of the classical genera of meristematic fungi. However, most species were described in the 19^th^ century, including the generic type species *C. olivaceum*, and voucher specimens are lost. Nevertheless, Haridas et al. (2020) introduced the family *Coniosporiaceae* and order *Coniosporales* using the few recently described rock-inhabiting fungi, *C. apollinis* and *C. uncinatum* as reference (Haridas et al. 2020). It should be noted that the family *Coniosporiaceae* was already used by Nannfeldt for the genus *Coniospora*.

### *Taxawa* gen. nov. Kurbessoian, Ahmed, de Hoog and Stajich -MycoBank (848163)

Type species: Taxawa tesnikishii

*Taxawa tesnikishii sp. nov.* Kurbessoian, Ahmed, de Hoog and Stajich -MycoBank (848164).

#### Description

Colonies moist, black, regular and flat on OA and heaped irregular on MEA and PDA, measuring 1.4-1.8 cm in diameter growing on MEA, OA, and PDA at 25°C. Hyphae smooth, thick-walled, measuring 6.1-10.3 μm wide. Yeast cells abundant produce unilateral budding 2.7-4.8 μm, subsequently inflating and becoming crucially septate and melanized with age. Clusters of black meristematic bodies (9.0-56.8 μm) were observed. Sexual state unknown.

#### Holotype

**CBS (150074), and herbarium CBS H-25287**, preserved at the herbarium collection of the Westerdijk Fungal Biodiversity Institute, Utrecht, The Netherlands. Living culture ex-type CBS (150074).

#### Diagnosis

Strain JES_119 showed similar morphology to the cultural states of e.g. *Sydowia* and *Comminutispora*, and has been described in the asexual genus *Phaeotheca*. Thambugala et al. (2014) erected the family *Aureobasidiaceae* for fungi in this relationship, but also this name existed already, *Aureobasidiaceae* Ciferri (1958) (Thambugala et al. 2014), based on the same species, *Aureobasidium pullulans* (Ciferri 1958). Only a small number of extant species has been sequenced, and the genome trees show an even larger taxon sampling effect, but given our current inability to assign strain JES_119 to any genus, we introduce the genus *Taxawa*.

## Discussion

BSCs are an untapped resource of novel microorganisms, using non-traditional methods of culturing may be a method to uncover more unavailable species for sequencing. Using a pour plate method for culturing allows the microorganism to avoid complete oxygen saturation (or oxygen toxicity), and instead can grow in low to micro-aerophilic concentrations of oxygen (Stevens 1995). Collecting MCF using this method has proved to be efficient, as fast-growing fungi overtake the surface of plates, while MCF can tolerate lower oxygen concentrations, thus localizing to the bottom of plates. Preparing axenic cultures of MCF is then relatively easy and can also increase the rate of isolation as MCF tends to take two weeks to grow. A hypothesis about the facultative aerobic or microaerophilic nature of MCF could also be seen in the phenotype of these species. Morphological growth on surface of petri plates shows unusual furrowing and clumping growth (JES_119, JES_115), some strains (JES_112) grow underneath the surface of the plate which seem to be MCF adaptations to the high concentration of oxygen present on petri plates.

The high melanin content of MCF complicates efforts to extract high molecular weight (HMW) DNA for sequencing, however adjustments to standard fungal CTAB can dramatically improve the gDNA purity. In the case for these BSC MCF, an increase in phenol:chloroform and chloroform washes allow for melanin to be removed completely from the DNA samples. Another consideration to help concentrate more DNA is to incubate the DNA sample in −20°C overnight during the DNA precipitation step. Using long read sequencing technology (Oxford ONT) it is best to have a very high concentration of clean DNA to process and thus could be the limiting factor for these sequencing procedures. Using hybrid methods of sequencing (Illumina and Nanopore) provides scientists the proper backbone to develop *de-novo* assemblies (Stevens 1995; Khezri et al. 2021; Chen et al. 2020; Saud et al. 2021). It is vital to have as near complete assemblies of species as possible, this provides the scientific community with the best tools for proper genomic assessments. For future research endeavors, a second nanopore flow cell would be necessary to help build a better assembly for strain JES_112 (*Neophaeococcomyces mojaviensis*) as the telomere analysis failed on this first round of assembly. The other two strains had enough depth of coverage that allowed for different programs to optimize the assembly and have more closed telomere contigs.

Scientific literature describing novel MCF always note the clonal or asexual nature, describing “missing” sexual states and thus always being in a haploid state. The MAT locus is one of the pivotal regions in the fungal genome that provides more accurate ploidy and sexual states of fungi, and using this data to describe the three new MCF assisted in understanding the sexual states. JES_115 and JES_119 as the two *Dothideomycetes* representatives indicate heterothallic sexual states having only the MAT1-2 gene, *vs.* homothallic species having representatives of both MAT1-1 and MAT1-2 genes. Better genome assembly would help better describe the JES_112 MAT locus as there is the presence of MAT1-2 but also a portion of the MAT1-1 gene (as it is composed of two genes, MAT1-1-1 and MAT1-1-4). The high melanin content of MCF could prove to be difficult when testing for ploidy in the scientific “gold standard” method (flow cytometry). But future research would most benefit from having this method optimized to understand not only the new MCF described in this work but also for future MCF considerations. Possibly the sexual states have not yet been described or seen for MCFs and may require a particular environmental stimulus to produce the proper structures to reproduce. Possibly, other forms of propagation are found in other habitats. A prototypical example is *Auerobasidium pullulans* which shows meristematic growth on glass or rock but is a rapidly expanding yeast-like fungus in culture. There has also been an increase in literature describing clonal MCF having high hybridization rates that create pseudo but stable diploids without the process of recombination (Gostinčar et al. 2022). This was not detected in these three species described in this paper.

MCF morphology indicates usual slow growth (about two weeks) seen in most MCF described previously (Selbmann et al. 2014; Chowdhary et al. 2014; Hölker et al. 2004; Zhao et al. 2010; Kejžar et al. 2013). JES_119, as it develops over time, increases its melanin production and develops from a bright orange-yellow color to a more deeply pigmented black. The budding cells are unilateral and are uncommon in other described MCF, and it was evaluated as a previously undescribed genus and species in Class *Dothideomycetes* as *Taxawa tesnikishii.* Strain JES_115 has a phenotype that is commonly seen in MCF, but due to its complicated taxonomy history, the genus *Coniosporium* remains as a genus of doubtful identity. The species JES_112 is one of the few described *Neophaeococcomyces* available anywhere and thus an important contribution to the scientific community.

Finding MCF from extreme environments such as BSCs may point to harbored secondary metabolites either absorbed from surrounding species through horizontal gene transfer (HGT) or adapted from time. MCF are known to use secondary metabolites to help survive such extremes and are thus an untapped resource for the identification of novel metabolites. An example to be described by Dr. Erin Carr, *Neophaeococcomyces* sp, has been shown to have an obligate bacterial hitchhiker (Carr 2022). JES_115 and JES_119 both contain about 15 secondary metabolites detected by antiSMASH, which is a considerable number when compared to other fungi (Rateb and Ebel 2011). The roles of secondary metabolites in modern human industry applications are vast (Demain 2014) and thus would be important to test and compare across all MCF.

## Conclusions

Biological soil crusts are an untapped resource for novel microorganisms as evidenced by our work. The MCF we collected are incredibly slow growing and could be easily missed during culturing attempts. The method described here (and detailed in **Appendix 1**) improved the chances for isolation of these species. MCFs have been isolated from different types of extreme environments including BSC’s.

An interesting consideration for species JES_112 or *Neophaeococcomyces mojaviensis* is through personal communications with another MCF scientist, Erin Carr, who also isolated a species of *Neophaococcomyces*, named “Crusty” from a biological soil crust. Their species was associated with a pink *Methylobacterium* symbiont, which she calls “Light Pinky”and “Dark Pinky”. Her assessment of this symbiosis suggests the role of the symbiont is involved in aerobic anoxygenic photosynthesis and auxin production (Carr 2022). Through our multiple culturing attempts, there was no indication of a pink symbiont, but the possible bacteria signal within our secondary metabolite assessment could be attributed to a contamination from this bacterium. JES_112 will be the second known genome of the genus *Neophaeococcomyces* found in GenBank as JAPDRQ000000000.

Species JES_119 or *Taxawa tesnikishi* is the first isolated of this phenotype. This species was closer to *Aureobasidium*, yet phenotypically is very different from identified *Aureobasidium* species. Using fungal phenotype and phylogenomics, we were able to determine the microorganism is different from known *Aureobasidium* species and may be of a new genus, hence the naming *Taxawa* as its genus name. *Taxawa tesnikishi* is the first known genus and species, now available in Genbank as JAPDRO000000000.

Species JES_115 or *Coniosporium tulheliwenetii* was also collected from the BSC’s from Boyd Deep Canyon and was thus honored with a Cahuilla term used to describe BSC phenotypic features seen on the soil. The genus *Coniosporium* has historically been confusing, as the earliest strains were collected in the late 1800s. Initial collections of *Coniosporium* were placed in Chaetothyriales, while later similar MCF phenotypes were also grouped with *Coniosporium*, to be later considered as Dothideomycetes instead. A careful reorganization and assessment of the genus is desperately needed. Genome assembly is now available as JAPDRP000000000.

## Acknowledgments

This work was performed in part at the University of California Natural Reserve System (Boyd Deep Canyon Desert Research Center) Reserve DOI: (doi:10.21973/N3V66D) (UCNRS Information Manager 1965) and at Joshua Tree National Park under permits generated through the help and guidance of Nuttapon Pombubpa, permit numbers: JOTR-2015-SCI-0033, JOTR-2016-SCI-0036, JOTR-2017-SCI-0044, JOTR-2018-SCI-0035, JOTR-2019-SCI-0031. Many thanks to Nicole Pietrasiak for her extensive knowledge in biological soil crusts. Many thanks also to Joshua Tree Reserve members who helped facilitate the admission of our Joshua Tree sample, specifically Anna R Tegarden, Ann Hitchcock, and Jay Goodwin.

